# Transgenerational epigenetic inheritance increases trait variation but is not adaptive

**DOI:** 10.1101/2024.04.15.589575

**Authors:** René S. Shahmohamadloo, John M. Fryxell, Seth M. Rudman

**Affiliations:** School of Biological Sciences, Washington State University, Vancouver, WA, United States; Department of Integrative Biology, University of Guelph, Guelph, ON, Canada

**Keywords:** Transgenerational epigenetic inheritance, Epigenetics, Phenotypic plasticity, Maternal effects, Genetic variation, Daphnia

## Abstract

Understanding processes that can produce adaptive phenotypic shifts in response to rapid environmental change is critical to reducing biodiversity loss. The ubiquity of environmentally induced epigenetic marks has led to speculation that epigenetic inheritance could potentially enhance population persistence in response to environmental change. Yet, the magnitude and fitness consequences of epigenetic marks carried beyond maternal inheritance are largely unknown. Here, we tested how transgenerational epigenetic inheritance (TEI) shapes the phenotypic response of *Daphnia* clones to the environmental stressor *Microcystis*. We split individuals from each of eight genotypes into exposure and control treatments (F0 generation) and tracked the fitness of their descendants to the F3 generation. We found transgenerational epigenetic exposure to *Microcystis* led to reduced rates of survival and individual growth and no consistent effect on offspring production. Increase in trait variance in the F3 relative to F0 generations suggests potential for heritable bet hedging driven by TEI, which could impact population dynamics. Our findings are counter to the working hypothesis that TEI is a generally adaptive mechanism likely to prevent extinction for populations inhabiting rapidly changing environments.

**One sentence summary:** Transgenerational epigenetic inheritance in *Daphnia* exposed to *Microcystis* revealed negative fitness effects on survival and growth rates, challenging hypotheses of a general selective advantage.

## Main Text

Anthropogenic global change is projected to drive significant biodiversity losses this century ^1^, highlighting the need to understand the mechanisms and magnitudes of adaptive phenotypic responses to environmental change ^2,3^. While organisms do undergo rapid evolutionary adaptation in response to environmental shifts ^4–6^, environmentally induced phenotypic plasticity represents the most general and impactful mechanism ^7,8^ and allows organisms to adjust phenotypes in response to the conditions they experience ^9^. Yet, most mechanisms underlying plastic shifts do not produce heritable change, limiting the long-term fitness benefits of plasticity when environmental fluctuations are common ^10^.

Non-genetic mechanisms of inheritance, broadly classified as ‘intergenerational’ or ‘transgenerational’, represent distinct biological pathways ^11^ and could play an important role in the transmission of heritable phenotypic changes in response to environmental fluctuations ^12^. Intergenerational inheritance involves the transfer of traits from parent to offspring through mechanisms independent of inherited DNA modifications, such as the acquisition of epigenetic marks during *in utero* development in live birth species and non-epigenetic mechanisms like maternal resource provisioning ^11,13,14^. Conversely, transgenerational inheritance involves the transmission of epigenetic information across multiple generations that can persist even in the absence of the original environmental stimulus—known as transgenerational epigenetic inheritance (TEI). The most studied underlying mechanisms of TEI are differential patterns of DNA methylation, histone modifications, and the transmission of non-coding RNAs ^11,15,16^. Despite methodological advances leading to a better understanding of inherited epigenetic marks associated with a variety of stressors across taxa ^17–19^, the true impact of epigenetic modifications on organismal phenotypes and population-level responses remains largely unknown.

An array of competing hypotheses propose the general effects of TEI on organismal fitness may be adaptive ^12,16,20–22^, nonadaptive ^23–25^, or maladaptive ^26,27^. One enticing hypothesis is that TEI might confer significant fitness benefits in response to environmental fluctuations ^22,23^, particularly in organisms with shorter generation times ^28^. This supposition is based on the assumption of an ‘epigenetic advantage’ ^23,29,30^, which posits that TEI leads to differential gene expression and can cause phenotypic change that ultimately enhances individual fitness. This assumption stems from the central dogma of genetics: differential DNA methylation patterns or histone modifications influence gene regulation in a direction that enhances fitness that could conceivably improve population persistence in changing environments ^10,31–35^. Testing the veracity of these adaptive assumptions requires measurements made in the F3 generation or beyond ^11,15,16,26^ to disentangle inherited epigenetic modifications from parental and non-inherited changes. To date, the limited number of studies measuring the phenotypic effects of TEI in the F3 and later generations ^27,36^ makes drawing definitive conclusions about its effects on phenotypes and fitness tenuous.

Relatively little is known or empirically demonstrated about the conditions under which the evolution of adaptive TEI would be anticipated ^26^, and—perhaps controversially to some—it is hypothesized that transgenerational epigenetic effects may not consistently confer benefits in adaptive plasticity under challenging environments ^25,26,37^. Existing empirical work on the phenotypic effects of TEI primarily relies on correlational findings due to the complexity of epigenetic modifications and their varied impacts on organismal fitness ^38–40^. Despite efforts to discover causal relationships, distinguishing between adaptive, nonadaptive, and maladaptive epigenetic changes remains challenging, as some modifications may lack discernable physiological consequences or remain silent ^24,40,41^. Another potential outcome of TEI is an overall increase in phenotypic variance, which could result either through selection and be categorized as ‘heritable bet hedging’ ^26^, or as a result of cumulative stress ^42^. Empiricists have been urged to test whether TEI produces adaptive, nonadaptive, or maladaptive mean phenotypic shifts and to quantify effects of TEI on trait variances in response to ecologically relevant conditions ^26^. Measuring TEI effects on fitness-associated phenotypes and projecting impacts ^43^ on population dynamics is critical for identifying adaptive or maladaptive phenotypic responses and their potential impact on population persistence in fluctuating environments. Doing so requires documenting TEI effects on fitness, and translating any putative effects to population-level outcomes requires measuring a suite of phenotypes associated with ‘vital rates’ that are the basis for population projection models ^44^.

*Daphnia* (water fleas) have proven to be a useful model system to study TEI because they reproduce clonally and investigations conducted across generations within clonal lines are rarely confounded by genetic variation and rapid evolution ^22,35,45^. *Daphnia* exhibits visible and measurable phenotypic responses to environmental perturbations that are key to population-level responses, including alterations in morphology, survival, and reproductive strategies ^46^. Harmful algal blooms (HABs) of the cyanobacterium *Microcystis* are a prominent aquatic contaminant ^47^ that can have both lethal and sub-lethal effects on a wide-range of taxa ^48–50^, including *Daphnia* ^51–53^. Many *Daphnia* populations show considerable intraspecific genetic variation and evidence of adaptation to HABs ^51,54–56^. Given the frequent and predictable nature of HABs, *Daphnia*’s tolerance to this stressor aligns with scenarios under which adaptive TEI would be expected to evolve ^12,16,21,22,37,57^. Studies on *Daphnia* in response to *Microcystis* have documented intergenerational plasticity after one generation of exposure ^58–61^ and TEI of environmentally induced DNA methylation ^62^. Testing whether TEI induces phenotypic shifts that impact *Daphnia* fitness in response to HABs provides empirical insight into the adaptive, nonadaptive, or maladaptive role of epigenetic inheritance ^26^ in a model system that has considerable ecological and conservation importance as a potential remediator of HABs ^63^.

To systematically address key questions regarding the adaptive potential of TEI in response to environmental change, we empirically investigated the following questions: i) Does TEI influence mean phenotypes? ii) If TEI influences mean phenotypes, is the direction of phenotypic change primarily adaptive, nonadaptive, or maladaptive? And, iii) Does TEI influence the amount of phenotypic variation? To determine whether TEI influences fitness-associated traits and population-level dynamics in *Daphnia* we compared two F3 exposure groups: ‘cccm’ (i.e., no great-grandmaternal exposure to *Microcystis* in F0 followed by great-granddaughter exposure to *Microcystis* in F3) and ‘mccm’ (i.e., great-grandmaternal exposure to *Microcystis* in F0 followed by great-granddaughter exposure to *Microcystis* in F3) repeated across 8 unique *Daphnia* clones (Figure 1). We quantified the chronic effects of the toxigenic cyanobacterium *Microcystis* on their life-history traits (survival, body growth, number of neonates produced, eye size, and time to first brood) in the final generation (F3). Clonal replication allows for an assessment of the effects of TEI across multiple genetic backgrounds and enables investigation of variation in the direction and magnitude of TEI across genotypes. By employing fitness-associated phenotypes measured across these 8 distinct tests within a simple population matrix model, we test for the effects of TEI on vital rates. This direct investigation provides valuable insights into a potentially significant mechanism governing organismal responses to environmental change, while also addressing critical questions ^22,26^ regarding the adaptive, nonadaptive, or maladaptive nature of TEI’s influence on mean phenotypes and its impact on the variance of phenotypic traits.

**Figure 1.**
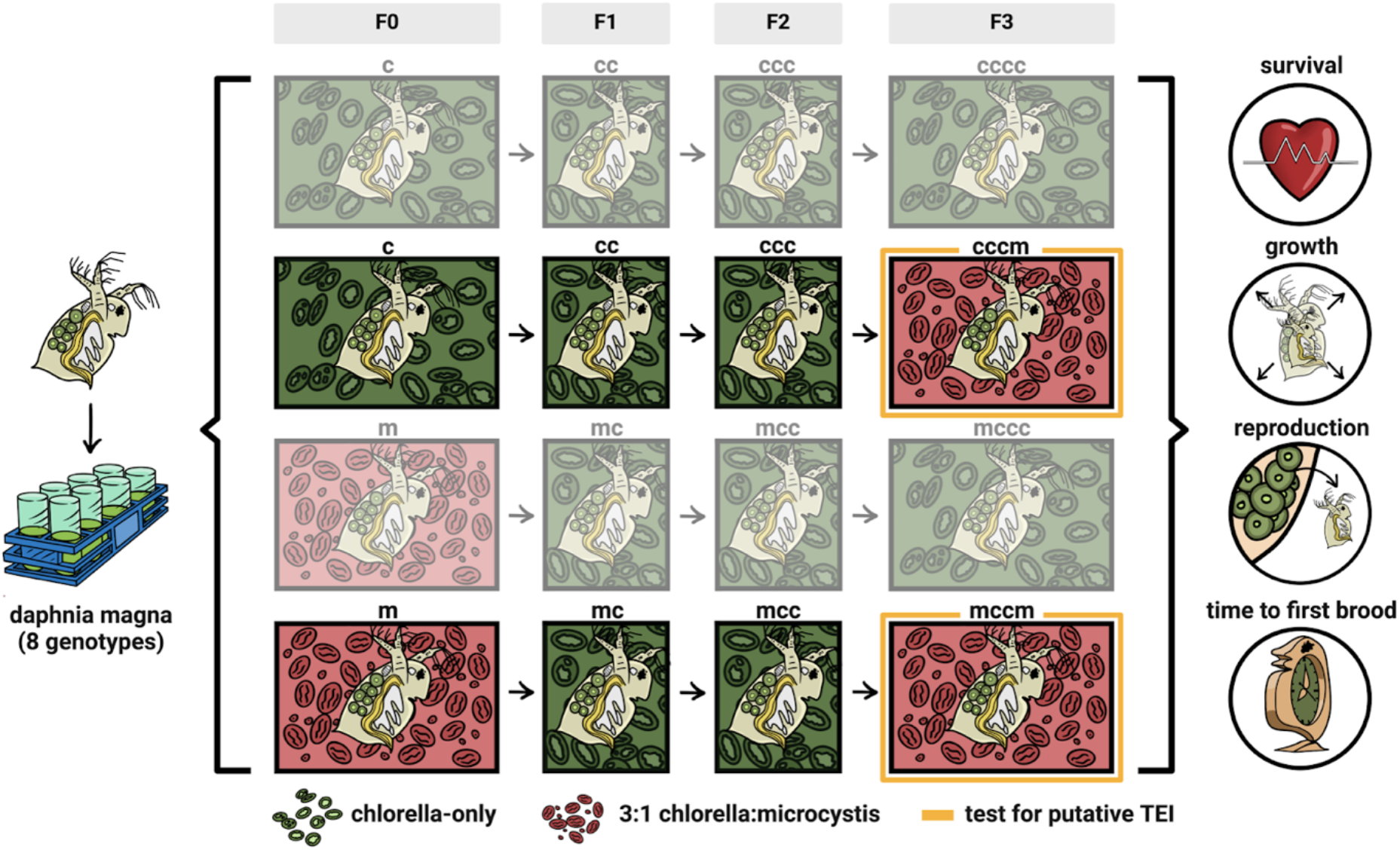
Experimental schematic to study transgenerational epigenetic inheritance. Eight genetically distinct clones of *Daphnia magna* were reared on either Chlorella-only (no stressor) or 3:1 Chlorella:Microcystis (stressor) in F0. Following this F0 treatment, F1 and F2 generations were all reared on Chlorella-only. To test for effects of transgenerational epigenetic inheritance, great-granddaughters (F3) across all populations were exposed to either Chlorella-only (‘cccm’) or 3:1 Chlorella:Microcystis (‘mccm’) (treatment series opaque with primary contrast highlighted in yellow). Relevant fitness traits were measured in this F3 generation to assess the impact of TEI and were used to create a population projection model.

## Results

### F0 – F2 generation impacts from exposure

All eight *Daphnia* genotypes had 100% survival up to their first reproductive event across all replicates in F0 → F2 when exposed to Chlorella-only (‘c’ → ‘ccc’). However, exposure to *Microcystis* (‘m’) caused a decrease in survival to an average of 67% across all *Daphnia* genotypes in F0, ranging from 57% survival in ‘genotype 5’ to 84% survival in ‘genotype 7’. Following cessation of *Microcystis* exposure, subsequent generations returned to near total survival to age at first brood (F1 (‘mc’) had 97% survival and F2 (‘mcc’) had 99% survival on average across all *Daphnia* genotypes (see Dataset S1, Supporting Information)).

### Great-grandmaternal exposure × F3 generation interactions

TEI, as measured on the F3 generation, caused a significant decrease in survival (χ^2^_(1)_ = 15.30, P < 0.01; Figure 2a). *Daphnia* from ‘cccm’ had survival rates of 78.75 ± 4.09% over 7 days compared to *Daphnia* from ‘mccm’ who had survival rates of 58.75 ± 6.11% survival over 7 days in F3. Similarly, we observed a significant delay in time to first brood (χ^2^_(1)_ = 16.08, P < 0.01; Figure 2d). *Daphnia* from ‘cccm’ reproduced at 11.03 ± 0.24 days compared to *Daphnia* from ‘mccm’ which reproduced at 12.55 ± 0.29 days in F3. A paired analysis showed a reduction in body size associated with TEI across all eight genotypes (t_65_ = −3.12, P < 0.01; Figure 2b), and the mean difference in body growth between ‘mccm’ and ‘cccm’ in F3 was estimated to be −14.15%. We did not observe significant effects of TEI on neonate production (χ^2^_(1)_ = 2.64, P = 0.10; Figure 2c). TEI did not produce detectable changes in eye size (χ^2^_(1)_ = 0, P = 0.99; Figure S1, Supporting Information), contrary to the hypothesis proposing maternal effects on offspring eye size as an adaptive response linked to improved foraging abilities ^61^.

**Figure 2.**
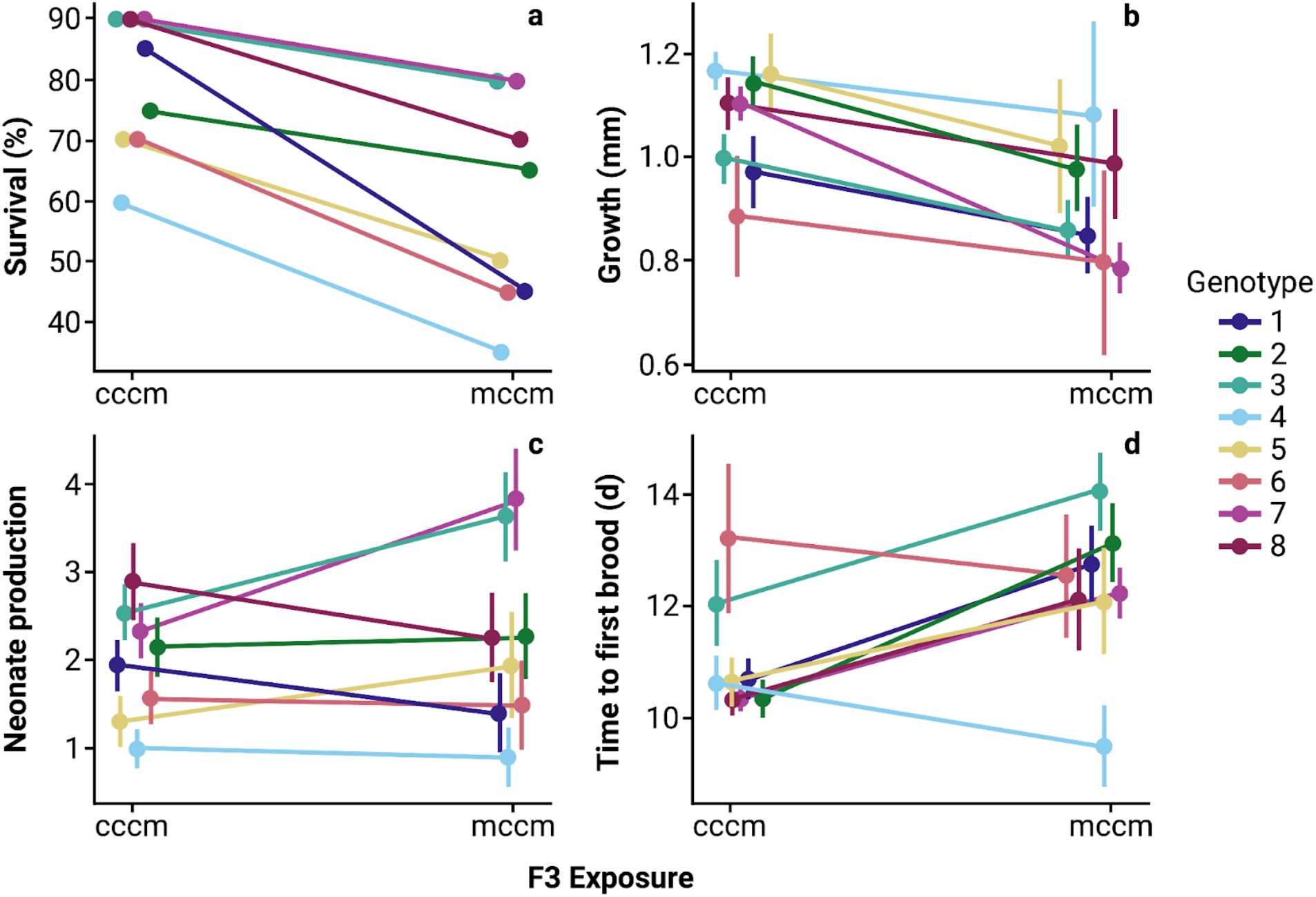
Phenotypic variation in a) survival at day 7, b) growth at day 7, c) neonate production, and d) time to first brood across eight *Daphnia magna* clonal populations after four generations (F0 → F3) of transgenerational epigenetic inheritance (±SE). *D. magna* exposed to *Microcystis aeruginosa* in F0 and F3 are signified by ‘mccm’, and *D. magna* only exposed in F3 are signified by ‘cccm’.

Survival reaction norms were additionally constructed to illustrate the range of survival rates exhibited by *Daphnia* under varying conditions of exposure to *Microcystis*, whether from great-grandmaternal exposure in F0 or great-granddaughter exposure in F3 (Figure S2, Supporting Information). Survival reaction norms depict the relationship between environmental conditions and an organism’s probability of survival, illustrating how survival rates vary across different contexts. Our results show that survival reaction norms were negative for all *Daphnia* genotypes in each exposure scenario (‘m’ compared to ‘c’, ‘cccm’ compared to ‘cccc’, ‘mccm’ compared to ‘mccc’) (Figure S2, Supporting Information).

### Great-grandmaternal exposure × population growth impacts

To assess the potential impact of TEI on population growth rates, we constructed Leslie matrices with observed mean rates of survival and neonate production from F3 exposure to ‘cccm’ and ‘mccm’. The net reproductive rate (R_0_) was next calculated for each F3 *Daphnia* clonal population, and the mean difference in R_0_ between ‘mccm’ and ‘cccm’ for each ‘clone’ and ‘exposure’ combination was measured to determine whether TEI exposure in F0 would be positive (R_0_ > 0), neutral (R_0_ = 0), or negative (R_0_ < 0) relative to populations with no mechanism for TEI (Figure 3a). TEI negatively impacted *Daphnia* clones 1 (R_0_ = −0.55), 4 (R_0_ = −0.10), and 8 (R_0_ = −0.65); TEI was neutral for *Daphnia* clone 6 (R_0_ = 0); and TEI was positive to varying degrees in *Daphnia* clones 2 (R_0_ = 0.15), 3 (R_0_ = 1.10), 5 (R_0_ = 0.65), and 7 (R_0_= 1.50). The mean difference in R_0_between ‘mccm’ (2.23) and ‘cccm’ (1.96) for all clones was 0.26, indicating that the overall demographic impact on *Daphnia* clones was neutral.

**Figure 3.**
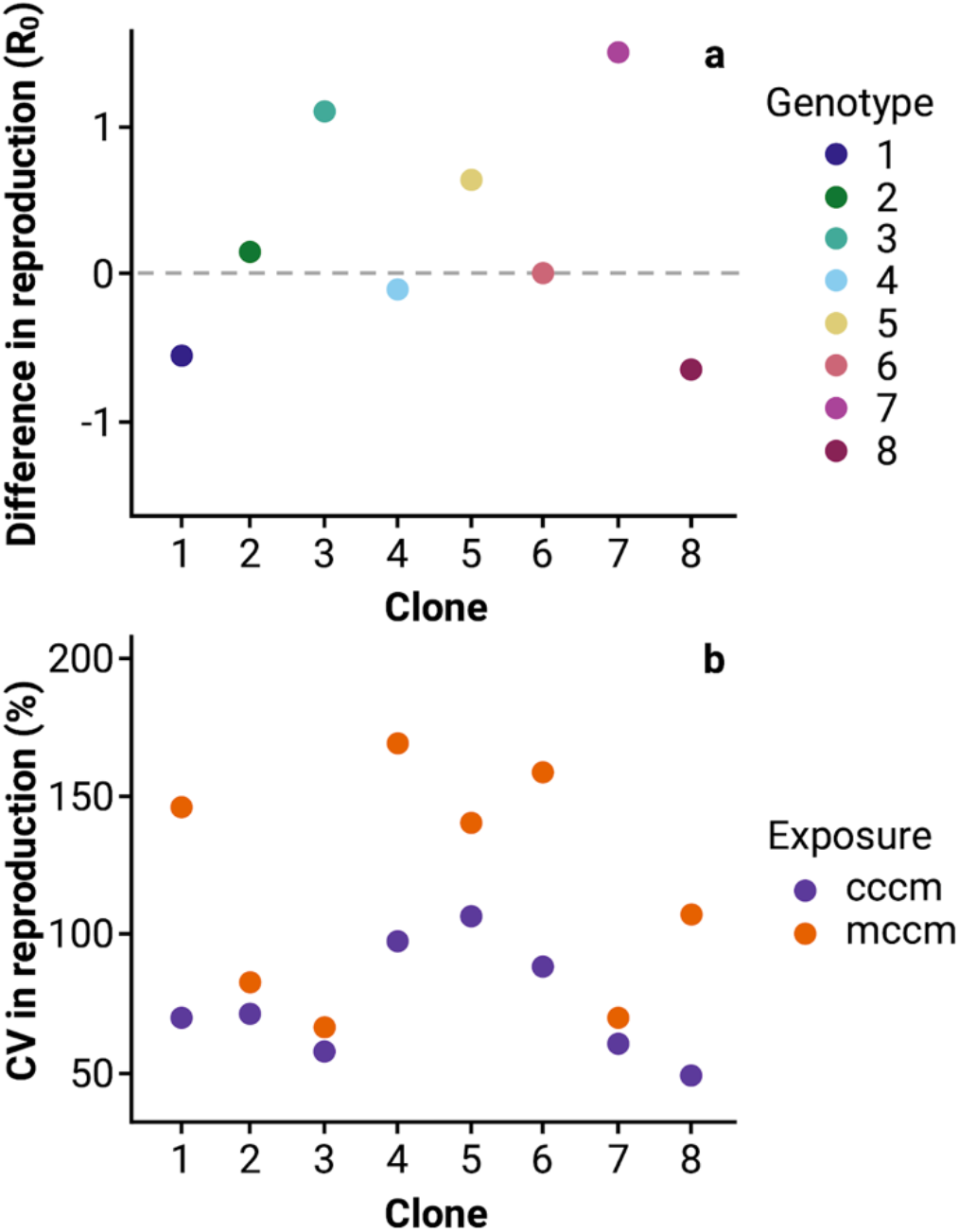
a) Difference in neonate production (‘mccm’ - ‘cccm’) between exposures to *Microcystis aeruginosa* across each of eight *Daphnia magna* clonal populations in the F3 generation. An adaptive response is >0 whereas a maladaptive response is <0. b) The coefficient of variation (CV) in neonate production of ‘mccm’ and ‘cccm’ exposures to *M. aeruginosa* across eight *D. magna* clones in the F3 generation.

Beyond shifts in trait means, changes in the variance of fitness associated traits can have profound impacts on populations ^64,65^ and can be a major driver of the pace of evolution by natural selection ^66,67^. Thus, we next calculated the difference in the coefficient of variation (CV) of neonate production between ‘mccm’ and ‘cccm’ from F3 exposure for each ‘clone’ and ‘exposure’ combination (Figure 3b). *Daphnia* with TEI exposure to *Microcystis* had significantly greater CVs than *Daphnia* from ‘cccm’ whose great-grandmothers were not exposed to *Microcystis* (F_1,14_ = 7.85, P = 0.014).

## Discussion

Overall patterns of phenotypic divergence associated with TEI exposure in F0 were considerable and the mean shifts we observed tended towards a maladaptive response across the 8 unique *Daphnia* genotypes (Figure 2) with evidence that TEI increases trait variance (Figure 3). When phenotypes were combined into a population projection, TEI did not lead to differences in population growth rates based on R_0_ (Figure 3). Given the ecological realism of the applied environmental stressor, which has been documented to induce substantial mortality ^47,52,55,68^, and prior work documenting notable transgenerational epigenetic modifications via DNA methylation in *Daphnia* exposed to *Microcystis* ^62^, the lack of adaptive TEI observed in our study is unlikely attributable to a weak environmental stressor.

Epigenetic mutations, if stable and beneficial, can significantly influence the rate and outcome of adaptation by speeding up the initial stages of “adaptive walks”, a progression wherein successive beneficial mutations drive a population closer to an optimal level of fitness ^21^. However, the impact of transgenerational epigenetic mutations on fitness values crucially depends on their stability, phenotypic effect, duration of the effect, and duration of the stressor ^16,69,70^. For TEI, if epigenetic mutations are unstable or have negative fitness effects, they may not persist across generations or may even hinder adaptive evolution ^21^. This theory runs contrary to other existing models suggesting the inheritance of acquired epigenetic variations can be adaptive across a wide range of environmental conditions ^71^ and can be beneficial in environments marked by predictable fluctuations ^22^. Recent population genetic models incorporating epigenetic variation further demonstrate the potential for stable epialleles to be maintained under neutral conditions and for epialleles compensating for deleterious mutations to deviate from mutation-selection balance, indicating a possible contribution of transient epigenetic regulation to the maintenance of genetic and epigenetic variation in populations ^72^. The latter theories are supported by recent experimental work in clonal yeast populations demonstrating that epigenetic switching, despite its instability, has adaptive advantages under particular fluctuating environments and can persist at low frequencies even in conditions predicted to be detrimental to epigenetic switchers ^73^.

With short generation times and inhabiting environments marked by intense seasonal HABs of *Microcystis, Daphnia* fit several key criteria for the evolution of adaptive TEI. We observed intraspecific genetic variation across clones for adaptive TEI (genotypes 3 and 7), suggesting there is standing genetic variation on which selection for TEI could act ^66^. Yet, notable constraints remain which may ultimately limit the evolution of adaptive TEI and explain the overall lack of positive TEI effects on fitness we observed. Epigenetic marks, such as DNA methylation or histone modifications, may not persist long enough for selection to effectively act on them due to their instability ^22,74^. These marks can be reversible and dynamic, potentially erasing or modifying in response to environmental changes or cellular processes ^19,22,75^. Given this instability, rapid adaptation from standing genetic variation might ultimately produce larger fitness benefits. Cases of rapid adaptation include evolution of phenotypic plasticity and intergenerational epigenetic inheritance, prominent in *Daphnia* responses to *Microcystis* ^54,55,76^. Our results support this; across environmental conditions (Chlorella-only and 3:1 Chlorella:Microcystis (present work), but also more severe HAB exposures (2:1 and 1:1 Chlorella:Microcystis ^56^), the 8 unique genotypes of *Daphnia* show both strong and consistent patterns of variation in fitness-associated phenotypes. Ultimately, the stability of epialleles, the frequency and predictability of environmental shifts, and the associated costs of epigenetic resetting via TEI, among other factors, may lead TEI to produce complex and unpredictable phenotypic outcomes ^26,77^.

Together these constraints may limit the frequency of adaptive TEI and, ultimately, support hypotheses that environmentally induced epigenetic changes are rarely truly transgenerationally inherited, let alone adaptive ^78^.

A notable directional effect of TEI on fitness associated phenotypes we observed was an increase in phenotypic variation—all 8 *Daphnia* clones had higher variation in F3 reproductive output with prior F0 exposure (Figure 3). This result could fit at least two potential mechanisms: 1) heritable bet-hedging (adaptive) or 2) increased variance due to cumulative stress (not adaptive). Heritable bet hedging describes cases where increased phenotypic variability provides a hedge against unpredictable environmental changes, increasing the likelihood of population persistence under fluctuating conditions ^26^. In contrast, the cumulative stress hypothesis ^42^ suggests repeated stressors could induce transgenerational effects on genome regulation that are maladaptive. Recent observations of compounded epigenetic impacts and disease susceptibility from successive multigenerational exposure to different toxicants in rats ^79^ demonstrates epigenetic modifications associated with cumulative stress. In line with the developmental system perspective ^80^, it is critical to consider the potential for TEI to arise from genome responses that mitigate short-term losses at the expense of long-term fitness effects, highlighting a more nuanced relationship between developmental plasticity, genetic mechanisms, and environmental change in shaping population dynamics.

The way in which increases in phenotypic variance influence the fate of populations dictates how these data should be interpreted. Increases in variation could have important and direct effects on populations, such as those described by Jensen’s inequality ^64^, or they could simply be maladaptive and lead to demographic costs. Over longer durations reliance on bet hedging strategies may result in extinction due to directional environmental changes ^26^, particularly in cases where stabilizing selection maintains a narrow range of trait values ^37^. More empirical data is needed to understand whether TEI generally increases phenotypic variance. Supplemented by further empirical investigations that measure or model the interaction between increased trait variation and varying amounts—as well as periodicity—of environmental variation, this approach could reveal whether the observed patterns of increased variance resulting from TEI confer adaptive advantages and are potentially significant for the maintenance of biodiversity ^81^.

Our study tests an array of competing hypotheses regarding the fitness effects of TEI in response to environmental stress. TEI exposure of *Daphnia* to *Microcystis* in F0 did not yield significant adaptive changes in fitness-associated phenotypes, revealing a propensity for maladaptive responses across clones. The absence of discernible effects on population growth rates rejects the hypothesis that TEI enhances population-level responses by *Daphnia* to cyanobacteria exposure. The observed increase in trait variation suggests there may be interesting potential for heritable bet hedging, with higher variance capable of influencing population persistence under challenging conditions. Our study calls for the construction of TEI models that better reflect the nuanced interaction between environmental stress, epigenetic inheritance, and standing genetic variation to better understand the mechanisms by which organismal phenotypes respond to fluctuating and challenging environments. While empirical investigations into TEI across taxa will help elucidate its role in organismal fitness ^26^, a reevaluation of its importance in population-level responses to environmental fluctuations is warranted.

## Methods

### Daphnia magna field collection and culturing

Eight genotypes of *D. magna* were collected from ‘Langerodevijver’ (LRV; 50° 49’ 42.08’’, 04° 38’ 20.60’’), a large waterbody (surface area = 140,000 m^2^, max depth = 1 m) within the nature reserve of Doode Bemde, Vlaams-Brabant, Belgium ^82^. In previous work we generated whole genome sequences of these clones which show they are genetically distinct and that tolerance to cyanobacteria is not correlated with metrics of genomic wide divergence between them ^56^. Like many temperate freshwater ecosystems LRV has yearly seasonal *Microcystis* HABs and contains a large resident population of *D. magna* (Luc de Meester *person. comm*.). Parthenogenetic lines of each genotype were maintained for over five years in continuous cultures in UV-filtered dechlorinated municipal tap water containing 2 mg C L^−1^ of the green alga *Chlorella vulgaris* (strain CPCC 90; Canadian Phycological Culture Centre, Waterloo, ON, Canada). *C. vulgaris* was grown in COMBO medium ^83^.

### Microcystis aeruginosa culturing

Following our previously described method ^84^, *M. aeruginosa* (strain CPCC 300; Canadian Phycological Culture Centre, Waterloo, ON, Canada) was cultured in BG-11 media and kept in a growth chamber under axenic conditions with a fixed temperature of 21 ± 1 ºC, cool-white fluorescent light of 600 ± 15 lx, with a photoperiod of 16:8 h light:dark. The culture was grown for a minimum of one month before preparation for the transgenerational plasticity study. *M. aeruginosa* CPCC 300 produces microcystins-LR (CAS: 101043-37-2, C_49_H_74_N_10_O_12_) and its desmethylated form [D-Asp^3^]-microcystin-LR (CAS: 120011-66-7, C_48_H_72_N_10_O_12_), which occur widely in freshwater ecosystems ^47,85^ and are toxic to many zooplankton species.

To prepare *M. aeruginosa* for testing on *D. magna*, an aliquot of the stock was inoculated in 100% COMBO medium for two weeks prior to test initiation and cultured to a cell concentration of 1.2 ± 0.02 × 10^7^ cells mL^−1^. This medium was chosen because it supports the growth of algae and cyanobacteria and is non-toxic to zooplankton ^83^.

### Transgenerational study

We evaluated within- and across-generation responses to *M. aeruginosa* using eight genotypes of *D. magna*. Phenotypic responses measured include survival, body growth, reproduction (number of offspring produced), eye size, and time to first brood.

To prepare for this study, we isolated one adult female *D. magna* per genotype in separate 50-mL glass tubes inoculated with COMBO medium and *C. vulgaris* at 2 mg C L^−1^, and monitored them daily for reproduction. *D. magna* juveniles born within 24 h were collected from their respective genotypes and individually separated into 50-mL glass tubes as previously described, totalling 10 replicates per genotype and 80 tubes total. These 80 *D. magna* individuals representing eight genotypes were the founding mothers of the transgenerational study, F0. All *D. magna* were incubated under constant conditions (temperature of 21 ± 1 ºC, cool-white fluorescent light of 600 ± 15 lx, with a photoperiod of 16:8 h light:dark).

To run this study, we reared F0 *D. magna* in one of two common gardens: Chlorella-only (optimal diet) and 3:1 Chlorella:Microcystis (toxic diet). Both common gardens provided *D. magna* with 2 mg C L^−1^, corresponding to 3 × 10^6^ cells total and corroborates with previous literature exposing daphnids to dietary combinations of green algae and cyanobacteria ^52,55,86^. The 3:1 Chlorella:Microcystis treatment was additionally chosen because these ratios exist in the wild ^47,85^ and can cause sublethal, intergenerational effects in *D. magna* ^52,53^.

A minimum of 40 replicates per F0 *D. magna* genotype, per common garden were individually raised in 50-mL tubes and fed their respective diets 3 × per week until they produced their first broods. All offspring across treatments were then reared for two generations —F1 and F2— in Chlorella-only until they too produced their first broods. The F2 offspring were then split in half for the F3 generation. The first subset of individuals (>20) from each clone were exposed to Chlorella-only until their first brood was produced. The second subset (>20 individuals) were exposed to 3:1 Chlorella:Microcystis. This combination of treatments generated a minimum of 40 replicates per original *D. magna* genotype in generation F0. Our previous work showed the magnitude of intraspecific genetic variation in the survival, growth, reproduction, and time to first broods of clones was significantly influenced by the presence of *M. aeruginosa* ^56^. To ensure 40 replicates per F2 *D. magna* genotype would survive to the final generation of F3 before it was split in half, we maintained additional replicates for certain genotypes that were particularly sensitive to *M. aeruginosa* toxicity. We individually tracked each *D. magna* replicate from mother to its daughter (F_x_ to F_x+1_) and tracked each *D. magna* great-grandmother to its great-granddaughter (F0 to F3) across all genotypes and common gardens. In summary, the experiment required a minimum of 640 F0 *D. magna* and 2,560 *D. magna* raised across all 4 generations, spanning 100 days (Figure 1).

Since this was a semi-static test, solutions were renewed 3 × wk by transferring *D. magna* from old to new glass tubes, followed by supplying each *D. magna* with 3 × 10^6^ cells of food, corresponding with 2 mg C L^−1^. Survival, reproduction, and the timing of first brood were recorded daily. Growth and eye size (mm) for each replicate across genotypes and common gardens were also measured on days 0, 3, 7, and day of the first brood for F0 and F3 to assess for TEI impacts within and across genotypes and treatment effects. The study was incubated under 400–800 lx cool-white fluorescent light at 20 ± 1 °C with a 16:8 light:dark cycle. Water chemistry parameters were measured at initiation, solution changes, and termination of the test.

### Statistical analysis

Phenotypic responses were analyzed using generalized linear mixed models (GLMM) with ‘great-grandchild exposure’ treated as a fixed effect and ‘*Daphnia* clone’ treated as a random effect to test whether great-grandchild exposure to *M. aeruginosa* (i.e., ‘mccm’ versus ‘cccm’) resulted in significant phenotypic differences. We fit appropriate family functions for each GLMM. For survival data, we used a binomial distribution and logit link. For neonate production, growth to day 7, and time to first brood datasets, we used a Poisson GLMM and log-link function.

For inferences on population impacts from TEI, Leslie matrices were respectively constructed for survival and fertility for F3 exposure to ‘mccm’ versus ‘cccm’. The population growth rate (λ) and net reproductive rate (R_0_) based on the Euler-Lotka equation were additionally calculated to estimate the rate of fertility in F3 *D. magna* mothers that were exposed to ‘mccm’ and ‘cccm’. These calculations were constructed using our early life data on *D. magna* (birth to time of first brood). This abbreviated life table may be representative of life histories in natural populations under high rates of predation ^87– 89^ and corresponds to the average duration of HABs ^90^. The difference in neonate production between ‘mccm’ and ‘cccm’ from F3 exposure were calculated for each ‘clone’ and ‘exposure’ combination and plotted. The difference in variance of neonate production between ‘mccm’ and ‘cccm’ from F3 exposure were also calculated for each ‘clone’ and ‘exposure’ combination and plotted. Levene’s test was utilized to assess the homogeneity of variances in the CV values between the ‘mccm’ and ‘cccm’ F3 treatment groups of *D. magna* mothers to determine whether significant differences existed in their neonate production.

For all analyses the *p*-level significance cutoff was 0.05. All statistical analyses were completed in R version 4.2.2 ^91^.

## Acknowledgements

This work was supported by an NSERC Postdoctoral Fellowship (R.S.S.), a Liber Ero Postdoctoral Fellowship (R.S.S.), the National Institute of General Medicine of the National Institute of Health Award #1R35GM147264 (S.M.R.), and a Canada First Research Excellence Fund through the Food from Thought Program (J.M.F.). We thank T. Gabidulin for his talented artwork in Figure 1, W. Smith for assistance with data collection, and the Rudman Lab for their valuable comments.

## Supporting Information

Dataset S1, Figures

